# Laminar differences in responses to naturalistic texture in macaque V1 and V2

**DOI:** 10.1101/710426

**Authors:** Corey M Ziemba, Richard K Perez, Julia Pai, Luke E Hallum, Christopher Shooner, Jenna G Kelly, J Anthony Movshon

## Abstract

Most single units recorded from macaque V2 respond with higher firing rates to synthetic texture images containing “naturalistic” higher-order statistics than to spectrally matched “noise” images lacking these statistics. In contrast, few single units in V1 show this property. We explored how the strength and dynamics of response vary across the different layers of visual cortex by recording multiunit and gamma band activity evoked by brief presentations of naturalistic and noise images in V1 and V2 of anesthetized macaque monkeys. As previously reported, recordings in V2 showed consistently stronger responses to naturalistic texture than to spectrally matched noise. In contrast to single unit recordings, V1 multiunit activity showed some preference for images with naturalistic statistics, and in gamma band activity this preference was comparable across V1 and V2. Sensitivity to naturalistic image structure was strongest in the supragranular and infragranular layers of V1, but weak in granular layers, suggesting that it might reflect feedback from V2. Response timing was consistent with this idea. Visual responses appeared first in V1, followed by V2. Sensitivity to naturalistic texture emerged first in V2, followed by the supragranular and infragranular layers of V1, and finally in the granular layers of V1. Our results demonstrate laminar differences in the encoding of higher-order statistics of natural texture, and suggest that this sensitivity first arises in V2 and is fed back to modulate activity in V1.

**Significance Statement:** The circuit mechanisms responsible for visual representations of intermediate complexity are largely unknown. We used a well-validated set of synthetic texture stimuli to probe the temporal and laminar profile of sensitivity to the higher-order statistical structure of natural images. We found that this sensitivity emerges first and most strongly in V2 but soon after in V1. However, sensitivity in V1 is higher in the laminae (extragranular) and recording modalities (local field potential) most likely affected by V2 connections, suggesting a feedback origin. Our results show how sensitivity to naturalistic image structure emerges across time and circuitry in the early visual cortex.

## Introduction

The perception of complex visual images arises through the activity of neurons across a series of hierarchically organized areas of the visual cortex. Beginning in the primary visual cortex (V1), neurons represent relatively low level image features such as local orientation and spatial scale. Near the top of the hierarchy, neurons in the inferotemporal cortex become selective for higher level properties like those defining particular objects or faces. The transformations linking these two representations are achieved through computations in mid-level ventral stream areas such as V2 and V4. However, understanding these intermediate visual representations and linking them to the anatomical organization of the visual cortex has been a continuing challenge to the field of visual neuroscience.

Recent work suggests that selectivity to the higher-order correlations found in natural images emerges downstream of V1. Natural images contain orderly structure which creates strong correlations in the output of V1-like filters tuned to different orientations, spatial scales, and local positions. These correlations can be measured from photographs and used to generate artificial stimuli with the same statistics (Portilla and Simoncelli, 2000). V2 neurons respond more vigorously to these “naturalistic” textures than to spectrally matched “noise” images lacking higher-order structure, while V1 neurons do not (Freeman et al., 2013). These results suggest that the ability of neural responses to discriminate between naturalistic and noise images – a property we refer to as “naturalness sensitivity” – arises through the transformation of visual information between V1 and V2. However, it is unknown how this selectivity may relate to known functional anatomy in either area.

The internal circuitry of V1 has been well studied (Callaway, 1998). Input from the lateral geniculate nucleus primarily drives cortical layer 4c, where receptive fields are more monocular and less orientation-selective (Garey and Powell, 1971; Hubel and Wiesel, 1972; Hendrickson et al., 1978; Blasdel and Lund, 1983). The granular layers of V1 in turn project to other laminae within the cortical column where neurons represent more complex properties of the stimulus, such as motion direction (Hubel and Wiesel, 1968; Livingstone and Hubel, 1984; Fitzpatrick et al., 1985; Hawken, Parker, and Lund, 1988; Yoshioka et al., 1994; Yabuta and Callaway, 1998). Functional differences across laminae in V2 are less well studied.

Measuring multiunit activity, we found the highest naturalness sensitivity in V2, consistent with previous single unit studies (Freeman et al., 2013). Surprisingly, we also found naturalness sensitivity in V1, although less than in V2. Naturalness sensitivity varied across cortical layers. It was highest in V2’s granular and supragranular layers, where the onset of sensitivity began soon after the visual response. The strength of sensitivity was followed by V2 infragranular layers, then V1 supragranular and infragranular layers, and finally near-zero in the granular layers of V1. The selectivity and timing of responses across layers suggest that naturalness sensitivity first develops in V2, while being fed back to the supragranular and infragranular layers of V1.

## Materials and Methods

### Electrophysiology

We recorded from four anesthetized adult macaque monkeys (three *Macaca nemestrina* and one *Macaca fascicularis*). Our standard surgical procedures were previously described (Cavanaugh, Bair, and Movshon, 2002). We maintained anesthesia with infusion of sufentanil citrate (6–30 μg kg^−1^ h^−1^) and paralysis with infusion of Norcuron; (0.1 mg kg^−1^ h^−1^) in isotonic dextrose-Normosol solution. We monitored and maintained vital signs within normal ranges (heart rate, lung pressure, EEG, body temperature, urine volume and specific gravity, and end-tidal pCO_2_). The eyes were protected with gas-permeable contact lenses and vision was corrected with supplementary lenses chosen through direct ophthalmoscopy. After data collection, the animals were sacrificed with an overdose of sodium pentobarbital. All experimental procedures were conducted in compliance with US National Institutes of Health Guide for the Care and Use of Laboratory Animals and with the approval of the New York University Animal Welfare Committee.

We made an occipital craniotomy and durotomy that allowed visualization of part of the operculum and lunate sulcus. We then implanted 32-channel linear arrays (NeuroNexus) near the estimated location of the border between V1 and V2. Electrodes were either spaced 50 or 100 μm apart so that arrays spanned 1.5 mm or 3.1 mm, respectively. We recorded from 12 electrode penetrations across the 4 animals.

### Stimulus generation

We generated all stimuli using the texture synthesis procedure described in Portilla and Simoncelli (2000). We used 32 different grayscale photographs (320×320 pixels) of visual texture. Each photograph generated one family of naturalistic textures, which included 20 different statistically matched samples of visual texture. The stimulus generation procedure was as given by Freeman et al., 2013 and Ziemba et al., 2016.

### Stimulus presentation

We presented visual stimuli on a gamma-corrected CRT monitor (mean luminance, 33 cd/m^2^) at a resolution of 1,280 × 960 with a refresh rate of 120 Hz. Stimuli were presented using Expo software on an Apple Macintosh computer. We characterized basic receptive field properties (orientation, spatial frequency, and size selectivity) of the neural activity found on each contact. We then presented naturalistic textures and their spectrally matched noise counterparts from 32 different texture families. The stimuli were randomly presented for 100 ms and separated by 100 ms of mean luminance. In some cases, stimuli were presented for 75 ms and separated by 225 ms of mean luminance, but results did not differ between the two conditions. All stimuli were centered at the estimated receptive field center of neural activity along the array and presented within a suitably vignetted aperture with a diameter of 4 or 6 degrees of visual angle. Each texture family was presented 70 times on average across experiments, meaning each unique sample was presented 3 or 4 times.

### Definition of cortical laminae

Animals were perfused transcardially with heparinized 0.01 M phosphate buffered saline (PBS) followed by a mixture of 4% paraformaldehyde (PFA) and 0.125% glutaraldehyde in 0.1 M phosphate buffer (PB), pH 7.3. Blocks of tissue surrounding the recording sites were removed and postfixed overnight in 4% PFA before sectioning parasagittally at 50 μm with a vibratome. Sections were incubated in 1% sodium borohydride/0.1 PB and rinsed in 0.1 M PB. The sections including electrode tracks were identified and histochemically stained for cytochrome oxidase by incubating overnight (and until the satisfactory completion of the reaction) at 37°C in 0.05% diaminobenzidine, 0.03% cytochrome c, and 0.02% catalase in 0.1 M PB. Sections were then rinsed in 0.1 M PB, mounted on glass slides, dehydrated in a graded series of ethyl alcohol solutions, cleared with xylene, and coverslipped.

For each penetration, electrode tracks through V1 were reconstructed to identify the total span of cortex, boundaries of cortical layers, and the location relative to the V1/V2 border. The absolute depth occupied by each layer was divided by the total cortical depth (pial to white matter) along the same penetration to yield the relative cortical depth spanned by each layer. To differentiate the supra-, infra-, and granular layers of V2, a Nissl counterstain was added to the relevant tissue sections. These were similarly reconstructed to identify laminar boundaries in units of relative cortical depth. Our estimates of cortical laminae divide the visual areas into three groups, supra-granular (layers 1-3), granular (all subdivisions of layer 4) and infra-granular (layers 5 and 6) sections. Our estimations of visual response latency based on our laminar estimations agree with previous literature.

### Single unit analysis

All data have been previously reported (Freeman et al., 2013; Ziemba et al., 2016) and fully detailed methods and further analysis can be found in those articles. We lowered quartz-platinum-tungsten microelectrodes (Thomas Recording) into V1 and V2 of 13 anesthetized, paralyzed, adult macaque monkeys (two *Macaca nemestrina* and 11 *Macaca fascicularis*). During each penetration we noted the depth where we estimated the gray matter of V1 and V2 to begin and end, as well as the depth at which each isolated single unit was recorded. Afterwards we normalized each single unit depth by the estimated beginning and end of cortex to achieve our relative cortical depth measurement. For all quantitative analyses, we averaged spike counts within a 100 ms time window aligned to the response onset of each single unit. We computed discriminability directly on these spike counts in the same manner as for our multiunit analysis.

### Multiunit analysis

We filtered the raw voltage signal (sampled at 30kHz) using a bandpass 2nd order Butterworth filter (corner frequencies 300 Hz and 6 kHz). We squared this filtered signal to give an instantaneous measure of spike-band power (Shooner et al., 2015). We low pass filtered and down-sampled the signal to 1 kHz and binned samples into bins of 5 ms. The first 30 ms of each trial was used to calculate the average baseline activity per electrode. Naturalness sensitivity was taken as the discriminability of responses to naturalistic and noise texture stimuli, computed by taking the difference between the means and dividing it by the shared standard deviation (d’). Naturalness sensitivity was computed for every 5 ms time bin for analysis of temporal dynamics. The time-averaged naturalness sensitivity for each site was computed on the multiunit signal averaged from 50 ms to 200 ms after stimulus onset.

Naturalness sensitivity was computed separately for each of the 32 texture families and then averaged. For the analysis comparing single unit, multiunit, and gamma activity, naturalness sensitivity was averaged across the five families evoking the strongest sensitivity in single units (Freeman et al., 2013), as not all families were shown to all single units. All other analyses averaged naturalness sensitivity across all 32 texture families.

### Local field potential analysis

We filtered the raw voltage signal using a bandpass 2nd order Butterworth filter to obtain a gamma band signal (20 Hz to 100 Hz). We then down-sampled the signal to 1 kHz, squared it, and binned samples into 10 ms bins to obtain gamma-band power over time. Discriminability values were calculated from gamma band power averaged from 50 ms to 200 ms after stimulus onset for each texture family and then averaged across families.

## Results

We used a model of natural texture appearance to generate synthetic stimuli for testing neural sensitivity to the presence of naturalistic higher-order correlations in images (Portilla and Simoncelli, 2000). This model captures the joint and marginal statistics of a simulated bank of V1 simple and complex cells (Fig. 1A). The parameters of the model include V1-like properties such as second order correlations specifying the orientation and spatial frequency content of an image (its power spectrum; Fig. 1A, middle). Other parameters represent correlations of higher-order that capture naturalistic features beyond the power spectrum (Fig. 1A, right). By iteratively adjusting seed images of Gaussian white noise (Fig. 1B, left), we generated sets of images having the same statistics as the original natural texture image, and refer to these images as “naturalistic textures” (Fig. 1B, right). We also generated images that contained the same power spectrum as the original natural image but lacked the higher-order correlations, and refer to these images as spectrally matched “noise” images (Fig. 1B, middle). Several previous studies have shown that while V2 neurons are sensitive to higher-order correlations – and thus fire more vigorously to the presentation of naturalistic stimuli – V1 neurons are not (Freeman et al., 2013; Ziemba et al., 2016; Okazawa, et al., 2017; Ziemba et al., 2018).

**Figure 1:**
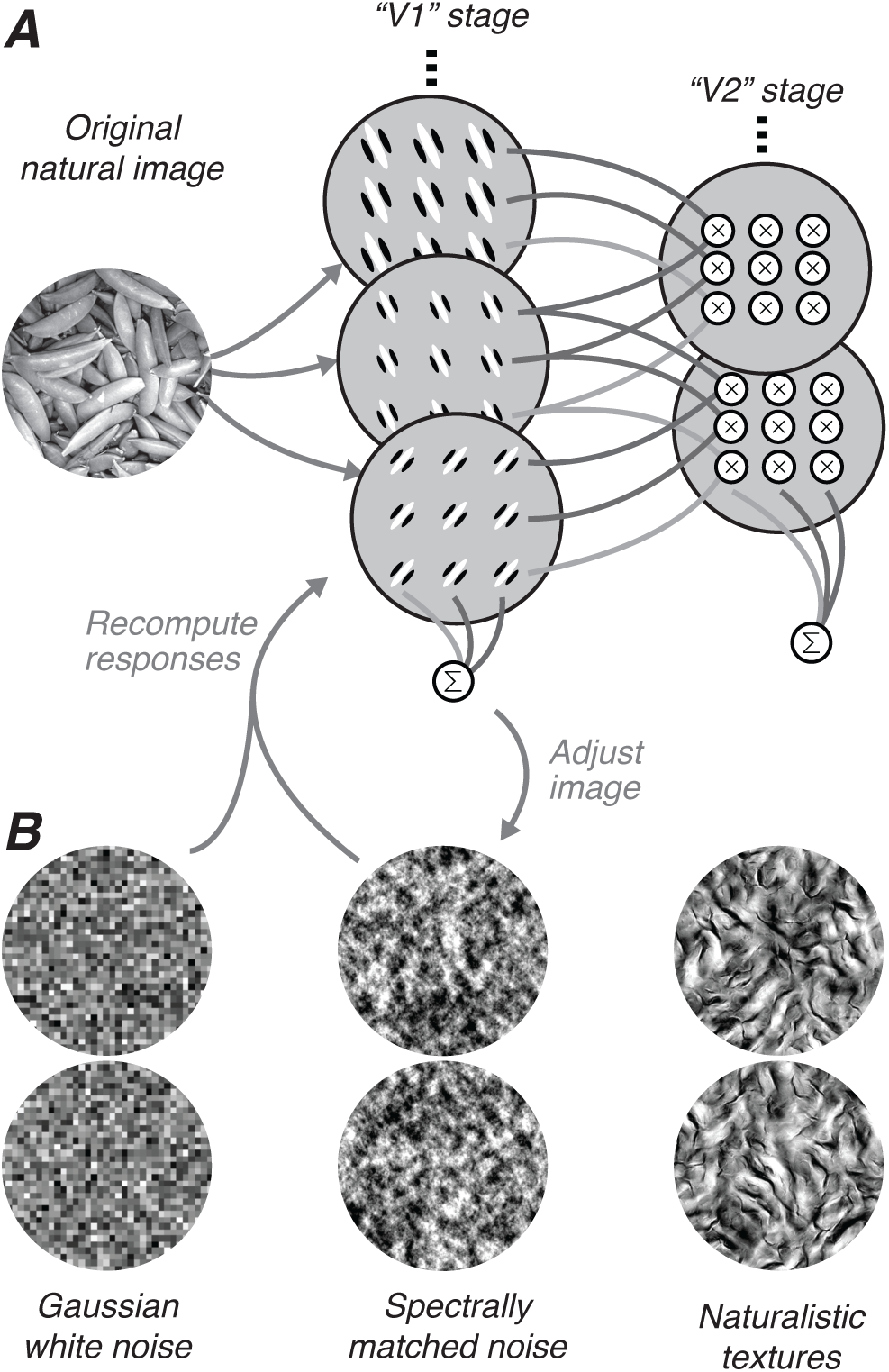
Stimulus generation. (A) Schematic of the model used for texture synthesis. An original black and white photograph (left) is decomposed into the responses of a population of V1 like filters tuned to different orientations and spatial frequencies (middle). The second stage computes local correlations (represented as products) across the output of filters tuned to different orientations, spatial frequencies, and positions. These correlations are then spatially averaged across all positions. (B) Schematic of the synthesis process. Each image is generated from a different seed image of Gaussian white noise (left). The white noise image is analyzed as in (A) and iteratively adjusted to more exactly match the statistics of the original image. If the image is matched for the full set of correlations, we refer to it as a naturalistic texture (right); if only matched for statistics capturing the spectral (second-order) content of the original image, we refer to it as spectrally matched noise (middle).

Previous work with naturalistic texture images randomly sampled single units from within a cortical area without reporting the laminar distribution. We reconstructed the relative cortical depth for a population of single units in V1 and V2 previously reported by Freeman et al., (2013). Sensitivity to naturalistic image structure has previously been captured with a modulation index (subtracting the response to naturalistic images from that to noise and dividing by their sum). We opted here to use signed d′ to facilitate comparisons to other recording modalities where the modulation index cannot be used. Greater d′ magnitude indicates that responses to the two types of stimuli were more discriminable, and a positive value indicates a larger response to naturalistic stimuli. For this population of V1 and V2 single units, signed d′ and modulation index were highly correlated (*r* = 0.75; *p* < 0.001). Although there did not appear to be much difference across layers in V1 or V2, the infragranular layers tended to yield stronger naturalness sensitivity in V1 (Fig. 2, left) and weaker naturalness sensitivity in V2 (Fig. 2, right). This is similar to a report in a study of V1 and V2 single unit sensitivity to multipoint pixel correlations (Yu et al., 2015). The data from that study and the data shown in Fig. 2 were recorded using independently movable electrodes, with the laminar localization determined offline. To provide a more reliable estimate of the relative cortical depth of naturalness sensitivity we decided to use linear multielectrode arrays in further experiments.

**Figure 2:**
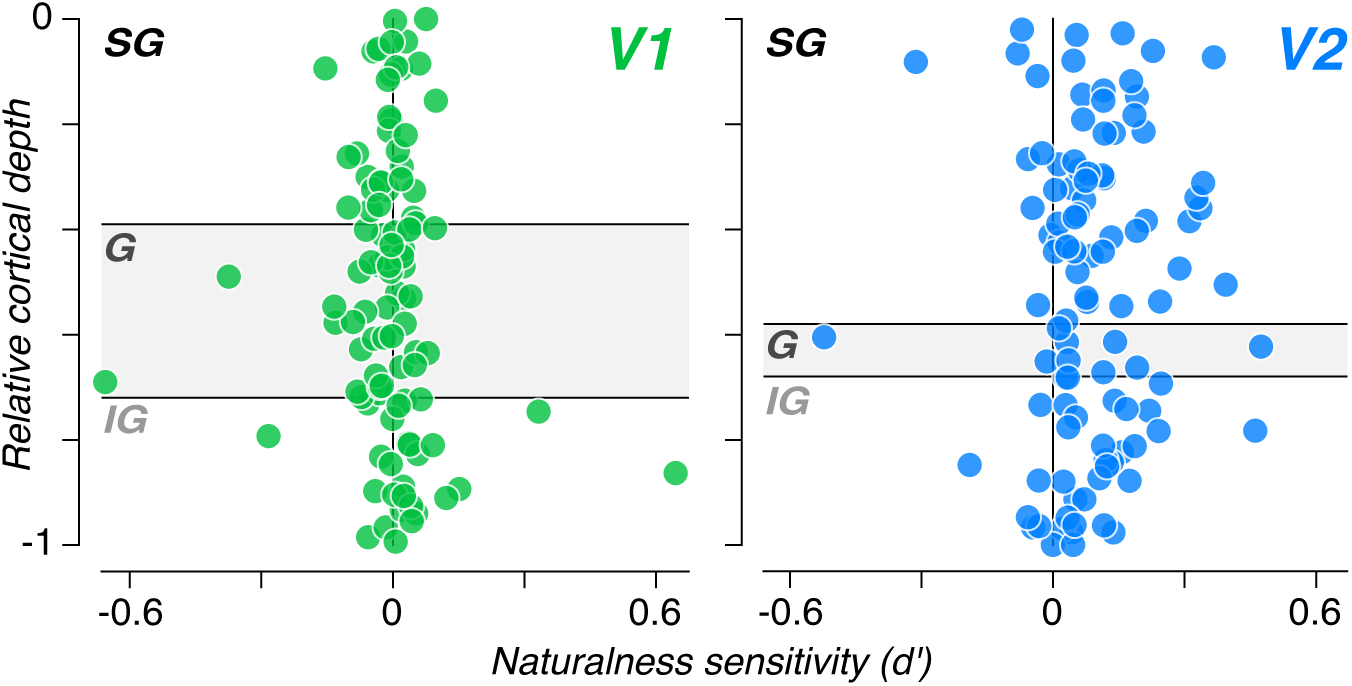
Cortical depth of naturalness sensitivity for single units in V1 and V2. Each circle represents one single unit from a population of 102 in V1 (left, green) and 103 in V2 (right, blue). Data were taken from Freeman et al. (2013). Laminar boundaries were drawn from tissue sections stained for NeuN and DAPI taken from Kelly and Hawken (2017), grouping layers into supragranular (SG), granular (G), and infragranular (IG) compartments.

We generally inserted our linear electrode arrays orthogonal to the cortical surface so that the electrodes spanned all cortical layers in either V1 or V2. Occasionally, we positioned the array to record simultaneously from both V1 and V2, and an example of such a penetration is shown in Fig. 3. Our electrodes spanned opercular V1, dorsal V2, and the intervening white matter (Fig. 3A). We confirmed this electrode placement offline histologically (Fig. 3A; See methods). We recorded strong, visually evoked multiunit activity from sites in both V1 and V2 in response to naturalistic and spectrally matched noise stimuli from a single texture family (Fig. 3B; there was little evoked activity recorded from electrodes in the white matter). The V1 sites generally showed similar multiunit activity to both naturalistic and spectrally matched noise, with some sites showing a late onset preference for naturalistic stimuli (Fig. 3B). In contrast, most V2 sites showed a strong preference for naturalistic stimuli over spectrally matched noise, and this preference emerged nearly simultaneously with response onset for most sites (Fig. 3B).

**Figure 3:**
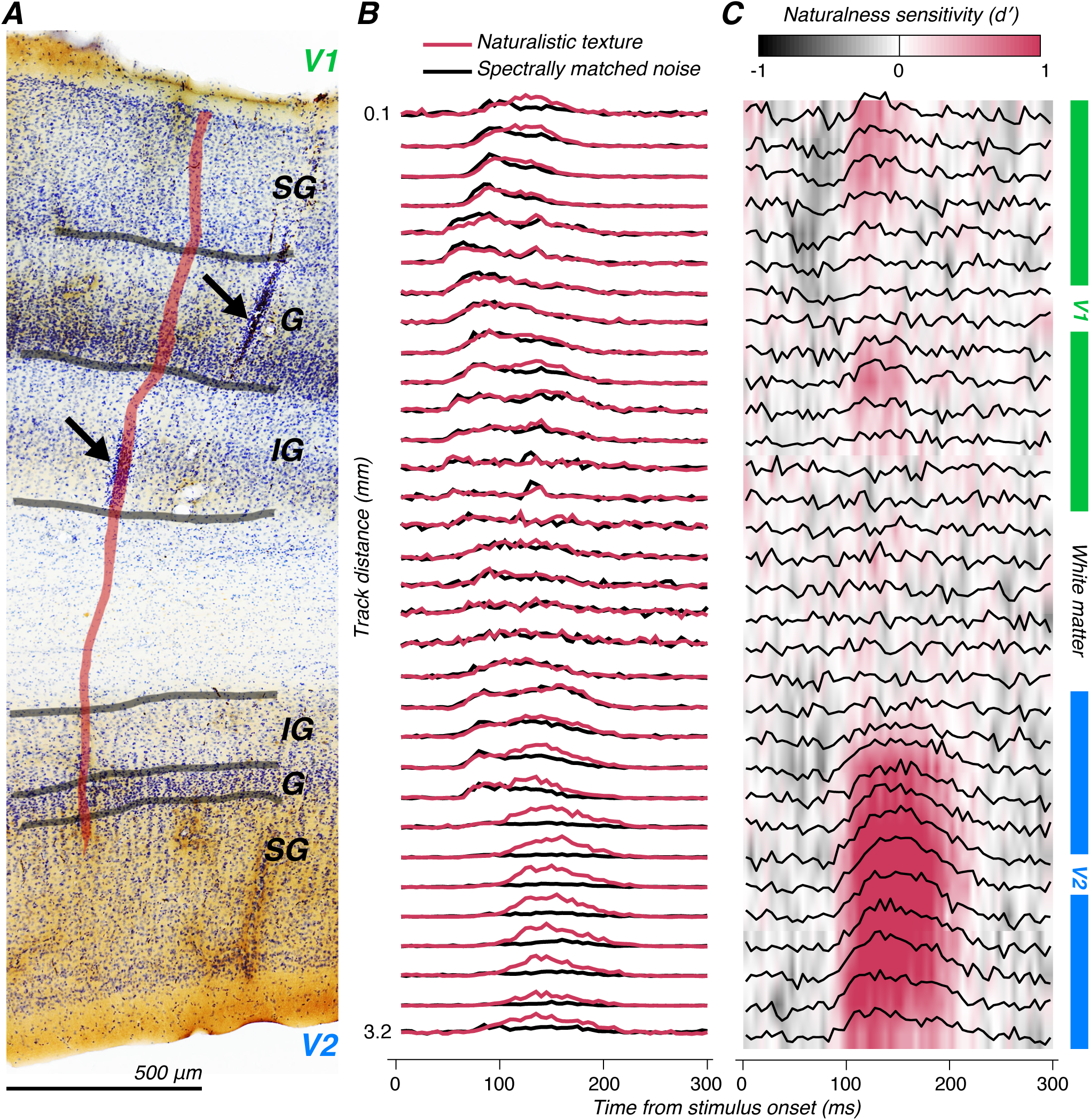
Example penetration. (A) Tissue section showing track damage from two laminar probe penetrations (arrows). The example penetration was identified across aligned sequential sections and followed a trajectory indicated by the red overlay. Laminar and gray/white matter boundaries were identified by staining for cytochrome oxidase and counter-staining for Nissl substance, and layers were grouped into supragranular (SG), granular (G), and infragranular (IG) compartments for analysis; these boundaries are indicated by gray contours. (B) Normalized multiunit responses to naturalistic texture (red) and spectrally matched noise (black) images from one stimulus family along the electrodes of the penetration illustrated in (A). The first visual responses appear in the granular layers of V1 (traces near the top), but the sites that differ the most in response to naturalistic and spectrally matched noise images are recorded from V2 (traces at the bottom). (C) d⊠ between responses to naturalistic and spectrally matched noise images as a function of time from stimulus onset across all electrode sites. Red indicates a higher firing rate to naturalistic texture, and black indicates a higher firing rate to spectrally matched noise. In V1, there is little sensitivity to naturalistic image structure at the beginning of response onset, but sites in the extragranular (but not granular) layers show a increase in d⊠ around 100 ms after stimulus onset. In V2, nearly all sites show high d⊠ with an onset that is roughly matched to the onset of the visual response.

The pattern of d′ across electrodes reveals further differences between V1 and V2 (Fig. 3C). In V1, there were clear laminar differences in naturalness sensitivity, with sites in the supragranular and infragranular layers of V1 responding with positive d′ after a delay, and sites in the granular layers responding with d′ near zero throughout. In contrast, nearly all V2 sites showed a higher and more sustained value of d′, and there was little variation across layers (Fig. 3C).

This pattern of results was representative of our findings across multiple penetrations. We made 12 electrode penetrations across 4 animals and grouped electrode sites from every V1 and V2 penetration into infragranular, granular, and supragranular layers based on histological reconstruction of electrode tracks. We computed naturalness sensitivity for each site. We averaged multiunit activity from 50 to 200 ms after the onset of naturalistic or spectrally matched noise stimuli from a single texture family. We then computed d′ from these responses and averaged across texture families to obtain a single measure of naturalness sensitivity for each site. After assigning all sites to a layer, we examined the distribution of d′ within each subdivision (Fig. 4). There was a significant effect of layer in V1 (*P* < 0.001, one-way ANOVA). The strongest naturalness sensitivity was found in the infragranular and supragranular layers (Fig. 4A), whereas d′ was near zero for the granular layers. However, each layer in V1 had a mean d′ significantly above 0 (*P* < 0.001, *t*-test). In V2, d′ was greater than 0 in all layers as well (Fig. 4B; *P* < 0.005, *t*-test). However, the magnitude of d′ did not significantly differ across V2 layers (*P* > 0.4, one-way ANOVA). Combining sites across all layers, naturalness sensitivity in V2 was significantly higher than in V1 (*P* < 0.001, *t*-test). The difference between V1 and V2 was not as great as previously found in single unit data (Fig. 2; Freeman et al., 2013), mostly as a result of higher naturalness sensitivity levels in V1. We wondered whether the apparent discrepancy with previous work might be due to our analysis of multiunit signals here in contrast to the previous focus on single units (Fig. 2; Freeman et al., 2013).

**Figure 4:**
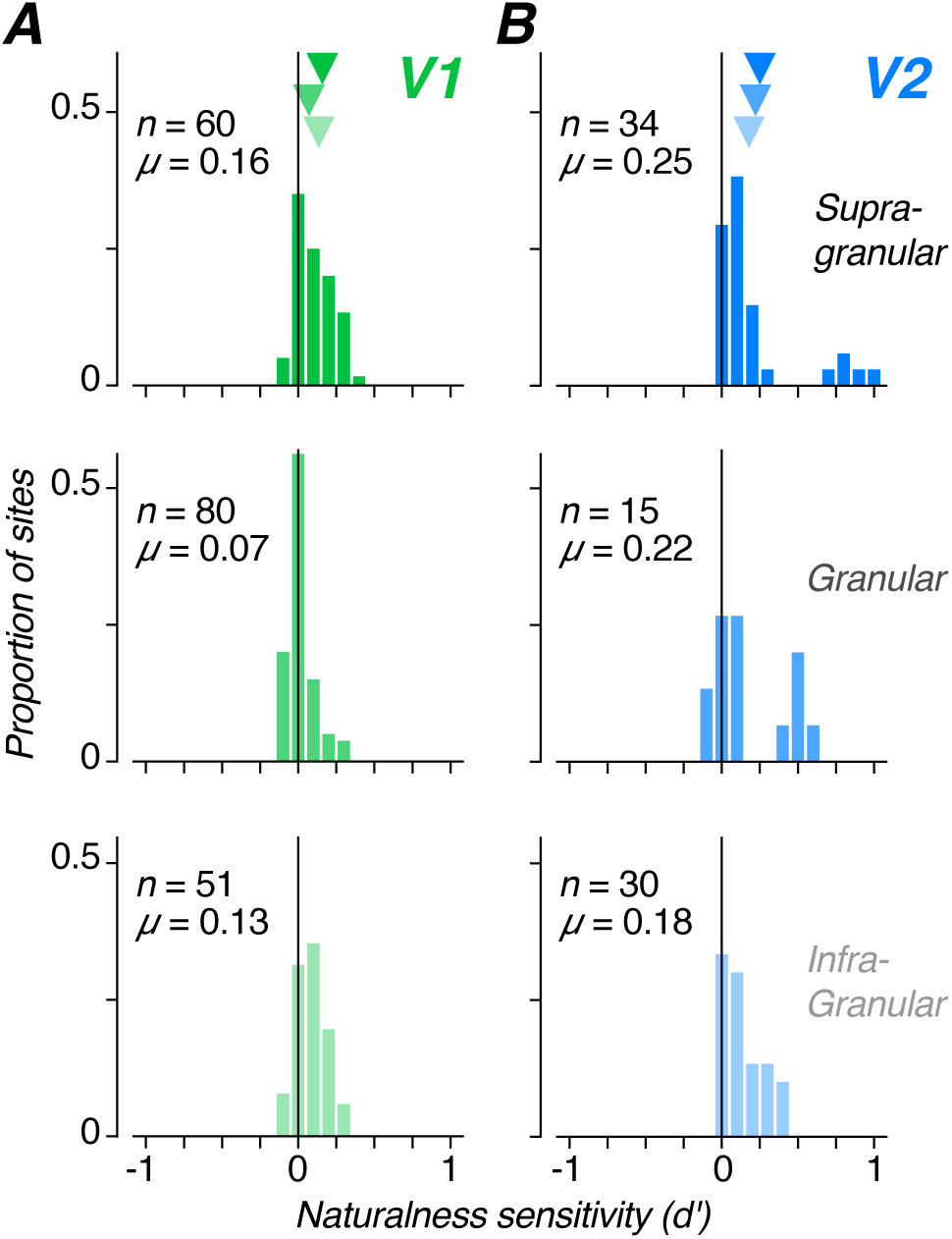
Distribution of naturalness sensitivity across layers. (A) Distribution of d⊠ between responses to naturalistic and spectrally matched noise images recorded from the supragranular, granular, and infragranular layers of V1. The mean value of d⊠ is indicated for each panel by the arrows in the top panel. (B) Same as (A) for sites located in V2.

To examine the effect of the neural signal modality on the strength of naturalness sensitivity, we reanalyzed previous single unit data in terms of d′ (Fig. 5A; data from Freeman et al., 2013). These data show very little naturalness sensitivity in the population of V1 single units (p > 0.05, *t*-test), but a slight bias towards positive values, while V2 single units are quite sensitive to the presence of higher-order statistics in naturalistic textures (Fig. 5A; p < 0.001, *t*-test). Compared with single unit responses, multiunit activity from the current experiments collapsed across layers showed more naturalness sensitivity in V2, and substantially more in V1 (Fig. 5B). However, V2 was still significantly more sensitive to the presence of naturalistic statistics than V1 (p < 0.001, *t*-test).

**Figure 5:**
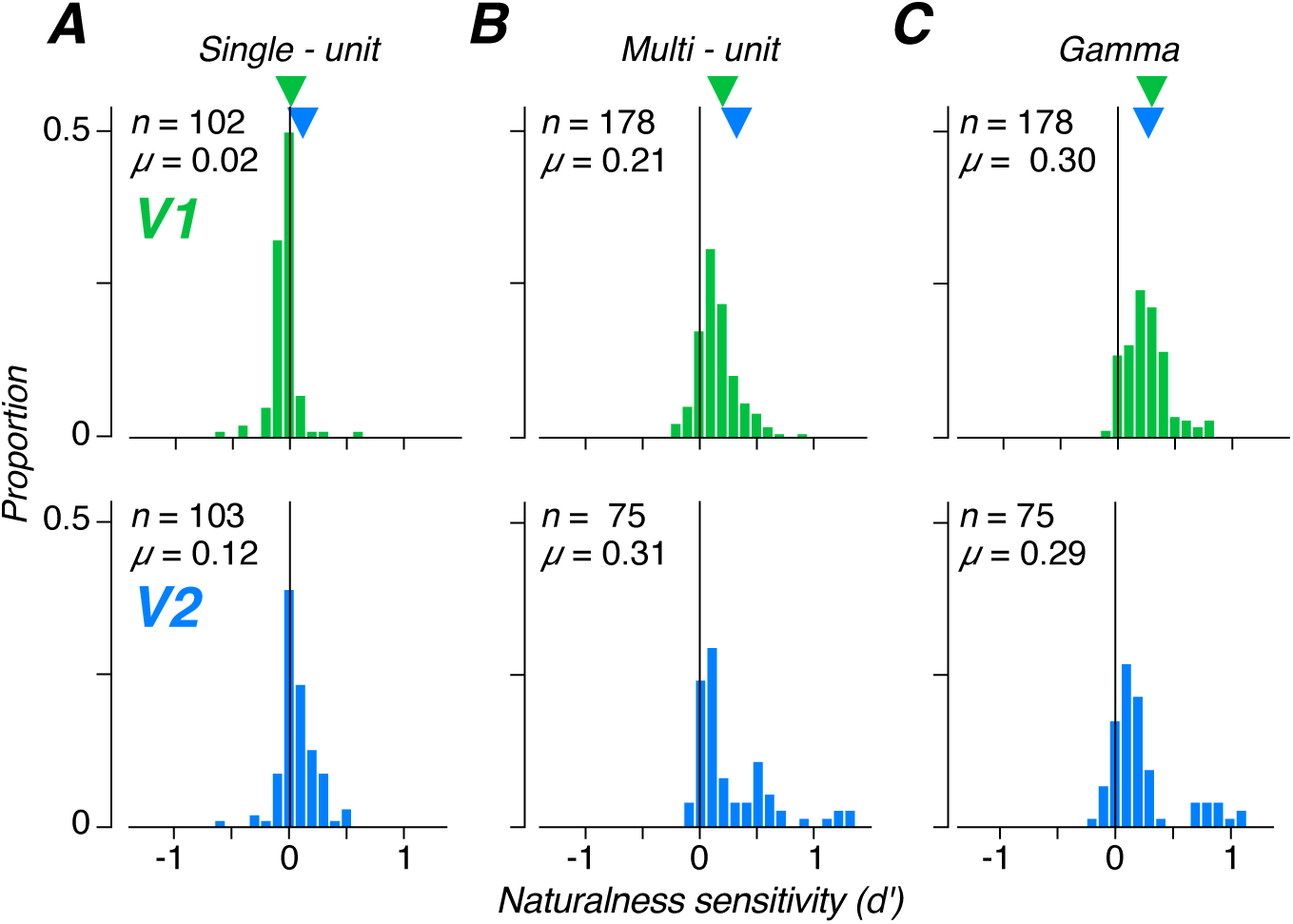
Distribution of naturalness sensitivity across recording modalities. (A) Distribution of d⊠ values computed from spike trains of single units recorded from V1 (top) and V2 (bottom). Data were taken from (Freeman et al., 2013). (B) Distribution of d⊠ values computed from multiunit responses in V1 (top) and V2 (bottom). (C) Distribution of d⊠ values computed from Gamma-band power (20 – 100 Hz) of the LFP in V1 (top) and V2 (bottom). The mean value of d⊠ is indicated for top and bottom panels by the arrows in the top panel.

We wondered whether the discrepancy between signal modalities in V1 could be explained by the tendency for the multiunit signal to be more influenced by sub-threshhold influences on membrane potential. To test this idea, we analyzed the gamma band (20-100 Hz) of the local field potential (LFP). The LFP signal is influenced by neural activity over a much broader area and is thought to be a reflection of the summed activity of subthreshold membrane activity (Katzner et al., 2009; Kang et al, 2010). In V1, LFP activity increases in the gamma band during visual stimulation, and its power may reflect feedback signals from higher visual areas (Kang et al, 2010). We found that the distribution of d′ was very similar in V1 and V2 when measured from gamma activity (Fig. 5C; p > 0.05, *t*-test).

Overall, V2 naturalness sensitivity was relatively stable across all signal modalities (Fig. 5, bottom). In contrast, V1 naturalness sensitivity increased from near zero with single unit spiking to approximately matched to V2 for gamma (Fig. 5, top). This pattern suggests that the sensitivity in V1 to higher-order statistics in naturalistic textures may be most prominently reflected in subthreshold membrane potential activity that could be the product of feedback connections from V2, V4, or higher visual areas where naturalness sensitivity is much stronger and is prominent in spiking responses (Okazawa et al., 2015, 2017; Ziemba et al., 2018). This feedback interpretation is also consistent with our laminar results. We recorded the strongest naturalness sensitivity in V1 from sites in the supragranular and infragranular layers, which are known to receive feedback from the supragranular layers of V2 (Fig. 4A; Mignard and Malpeli, 1991). We observed the weakest naturalness sensitivity from sites in the granular layers of V1 which do not receive feedback inputs. A plausible account of our results thus suggests that sensitivity to higher-order statistics emerges through computations performed by V2 neurons on their V1 inputs, and is subsequently fed back from V2 to modulate V1 responses – manifesting weakly in spiking responses, and strongly in signals that reflect subthreshold membrane fluctuations.

To explore this idea, we examined the temporal dynamics of multiunit responses in V1 and V2. We normalized all multiunit responses to their maximum and averaged all sites as a function of time. The early portion of the visual response in all layers did not appear to differ between naturalistic and spectrally matched noise stimuli in V1 (Fig. 6A, red and black lines). We computed average d′ as a function of time in each set of layers and found that it did not deviate from zero until tens of milliseconds after the onset of the visually driven response (Fig. 6A, green line). When we performed the same analysis in V2, we found a different pattern. Naturalness sensitivity in V2 (Fig. 6B, blue line) emerged soon after visual response onset (Fig. 6B, red and black lines), particularly in the supragranular and granular layers.

**Figure 6:**
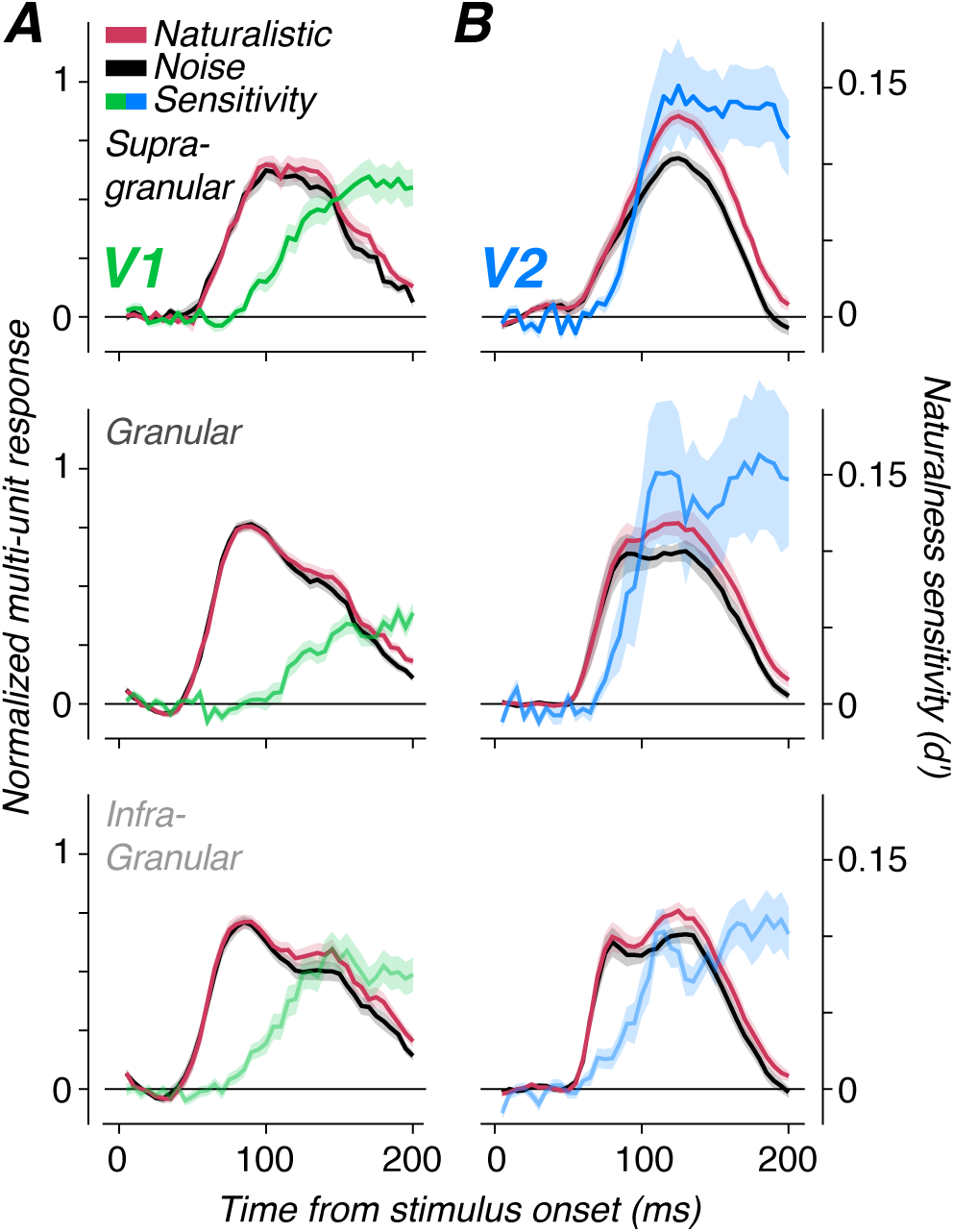
Response dynamics across layers. (A) Average V1 multiunit response to naturalistic (red) and spectrally matched noise (black) stimuli as well as the d⊠ between them (green) as a function of time from stimulus onset for supragranular, granular, and infragranular cortical layers. The ordinate axis for the normalized multiunit response is shown on the left, and the ordinate axis for naturalness sensitivity (d⊠) is shown on the right in panel (B). (B) Same as (A) for sites located in V2. Shaded regions indicate ±s.e.m.

To examine these effects more directly, we plotted the average time course of naturalness sensitivity for all layers in both V1 and V2 (Fig. 7B) as well as the cumulative d′ (Fig. 7D). Sensitivity in all layers of V2 was always higher than in all layers of V1 throughout the early phase of the response and appeared to have an earlier onset. Naturalness sensitivity began to rise across layers in V2 between 70 and 80 ms post stimulus, whereas naturalness sensitivity was absent in V1 until roughly 10 ms later when the supragranular and infragranular layers of V1 began to discriminate naturalistic and noise stimuli (Fig. 7B,D). The granular layers of V1 were even further delayed, with no naturalness sensitivity evident until after 100 ms post-stimulus.

**Figure 7:**
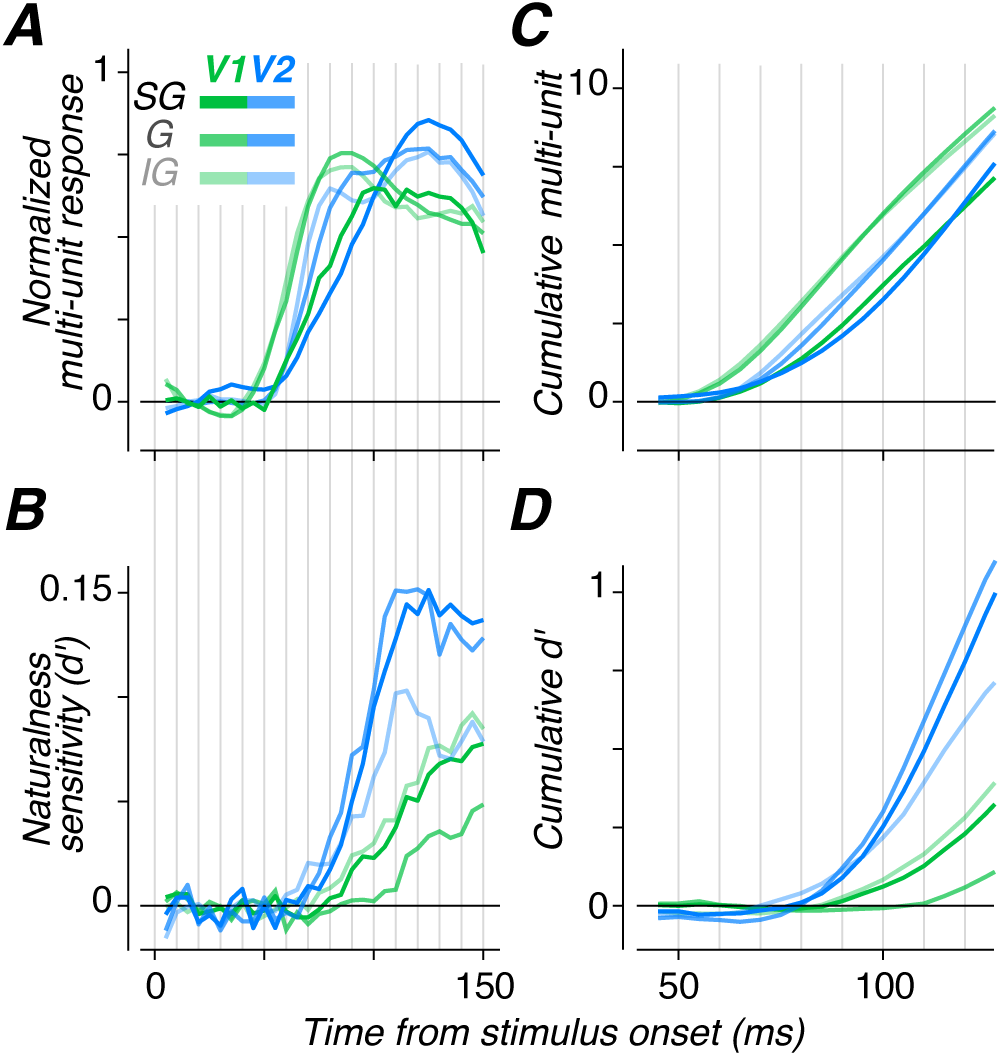
Dynamics of visual response and naturalness sensitivity differ across layer and area. (A) Average multiunit response to naturalistic texture stimuli as a function of time from stimulus onset for different layers (indicated by shading; SG: Supragranular; G: Granular; IG: Infragranular) in V1 (green) and V2 (blue). (B) Average d⊠ between naturalistic and noise as a function of time from stimulus onset for different layers (indicated by shading) in V1 (green) and V2 (blue). (C) Cumulative of the average multiunit response shown in (A). (D) Cumulative of the average d⊠ shown in (B).

The timing of the onset of naturalness sensitivity is quite different from the onset of visual responses (Fig. 7A,C). We examined the dynamics of multiunit activity in response to naturalistic images and found the expected pattern of visual response across layers. The granular and infragranular layers of V1 began to respond first, followed by the granular and infragranular layers of V2, and finally the supragranular layers of V1 and V2 began responding (Fig. 7A,C). In contrast to this flow of visual information, naturalness sensitivity appeared to emerge through computations performed in V2 and only subsequently appeared in the V1 multiunit response of the supragranular and infragranular layers, presumably because of the delayed influence of feedback connections.

## Discussion

We investigated the emergence of sensitivity to the statistics of natural textures by examining neural activity across stages of cortical processing in V1 and V2. Compared with responses in V1, V2 sensitivity for naturalistic image structure was stronger and emerged earlier. In V1, the supragranular and infragranular layers exhibited more naturalness sensitivity than previously reported for single unit recordings, although still less than in V2. Since these layers of V1 receive prominent projections from V2 (Rockland and Virga, 1989; Mignard and Malpeli, 1991), we suggest that naturalness sensitivity in V1 may be largely due to feedback. Furthering this point, V1 sensitivity emerged after that in V2 and was delayed relative to the onset of the V1 visual response. Thus, the pattern and dynamics across laminae suggest that sensitivity to the higher-order statistics of natural textures is achieved through computations first performed by neurons in V2. Sites in the supragranular and infragranular layers of V1 subsequently become weakly sensitive through the influence of feedback.

Our finding of significant sensitivity to higher-order correlations in V1 appears to contradict previous reports of its absence in the firing rate responses of V1 single units (Freeman et al., 2013). Here, rather than recording single unit spiking, we analyzed multiunit activity by measuring high frequency power in the LFP (Shooner et al., 2015). The multiunit signal is thought to incorporate activity from units within 30 to 100 µm of the electrode contact (Buzsaki, 2004; Xing et al., 2009), and thus aggregates many weak signals from individual units into a stronger differential signal between naturalistic and noise images. We demonstrate that signal modality can indeed have strong effects on the measured d′ by comparing the multiunit to LFP gamma band activity at the same sites (Fig. 5). Gamma is thought to reflect accumulated neural activity and postsynaptic potentials from a wide region of cortex surrounding the electrode contact (Katzner et al., 2009; Kang et al, 2010), and has been shown to differ in its selectivity from spiking responses under some circumstances (Jia et al., 2011; Berens et al, 2008). Thus, as the recorded signal incorporates information across a broader region of cortex and lower frequencies, V1 neural activity appears to become more sensitive to the presence of higher-order correlations in the stimulus. We posit that this may be because V1 multiunit and gamma activity are more affected by feedback from V2 or other extrastriate areas. Since the LFP reflects cumulative membrane potential fluctuations beyond spiking, it may be more reflective of top-down processes and the influence of feedback (Kang et al., 2010; Jia et al., 2011; Van Kerkoerle et al., 2014). Our multiunit signal may lie at an intermediate position between isolated single units and gamma on a spectrum of feedback integration from higher visual areas.

Previous evidence also points to the influence of feedback on V1 responses to naturalistic visual texture. Average V1 single unit responses exhibit a late sensitivity to higher-order correlations, about 80 ms after the onset of visual responses (Freeman et al., 2013). Although the naturalness sensitivity of the full visually evoked response in V1 was uncorrelated across different textures with that in V2, the naturalness sensitivity of this late V1 response component was significantly correlated with V2 sensitivity (Freeman, 2013). Additionally, although the BOLD response to alternation of naturalistic and noise stimuli was significantly higher in human V2 compared with V1, V1 exhibited significant modulation (Freeman et al., 2013). This weak V1 BOLD modulation was also highly correlated with V2 BOLD modulation and the V2 single unit modulation across different textures (Freeman et al., 2013; Freeman, 2013). This pattern of results suggests the influence of modulatory feedback, which is thought to be more prominent in BOLD fMRI than spiking signals (Ress et al., 2000).

There has been an ongoing debate about what components of neural activity are best represented by the BOLD signal. LFP can better predict BOLD than multiunit activity under some conditions (Logothetis, 2002), although the difference is not great and may depend on longer stimulus presentation times than used in our experiments (Heeger and Ress, 2002). Here, we find qualitative differences in the pattern of naturalness sensitivity in V1 and V2 across single units, multiunits, and LFP. Previous BOLD experiments are clearly best matched to the multiunit spiking analyzed here (Freeman et al., 2013) in that V1 sensitivity is greater than 0 but a substantial difference in sensitivity remains between V1 and V2. Our results are thus consistent with recent studies that find a better match between multiunit activity and BOLD than between LFP and BOLD (Lima et al., 2014), and the use of naturalistic texture may thus provide a tool for further studying the relationship between different recording modalities.

Recent studies have found similar laminar differences between V1 and V2 in the representation of contours embedded in a randomly oriented background (Chen et al., 2017). Like our results here, Chen et al. (2017) found the highest sensitivity to contours in the supragranular and granular layers of V2 and in the supragranular and infragranular layers of V1, and they also attribute this pattern of sensitivity in V1 largely to top-down feedback from V2. However, our results provide two distinct insights about the selectivity and dynamics of laminar circuitry in early visual cortex. First, the recordings in Chen et al. (2017) are made from monkeys trained to perform a contour grouping task, and so their results may result from strong top-down influences reflecting cognitive states such as attention (van Kerkoerle et al, 2017). Our experiments were performed under anesthesia, so represent only reflexive feedback mechanisms operating in the absence of goal-directed behavior. Second, sensitivity to embedded contours found in V2 is likely due to the larger receptive field sizes of V2 neurons (Chen et al., 2017). Thus, contour selectivity in V2 merely reflects integration of orientation signals within the classical receptive field, and not a more complex form of selectivity. In contrast, sensitivity to naturalistic image structure reflects neural computations distinctive to V2 (and not due to receptive field size differences with V1, Freeman et al., 2013), and a potential functional signature for investigating feedback to V1 without the need to tailor stimuli to the receptive fields under study.

Our results complement other recent work examining responses in V1 and V2 to synthetic binary texture stimuli containing particular multipoint correlations. Yu et al. (2015) found that sensitivity to multipoint correlations emerges most strongly in the supragranular and granular layers of V2, and although very weak in V1, appears stronger in the supragranular and infragranular layers compared with the granular layers. Additionally, they found that the correlations represented most prominently in V2 were those that were most informative when extracted from natural images (Hermundstad et al., 2014), suggesting that the higher-order correlations in our stimuli and theirs could result in similar image structure. Further work is needed to quantify the relationship between multipoint correlations and the higher-order image statistics used here, but both approaches capture information about a visual scene beyond the local power spectrum (Portilla and Simoncelli, 2000; Hermundstad et al., 2014). The similar pattern of laminar activity in these studies is also promising evidence that sensitivity to isolated higher-order statistics of binary texture patterns may reflect the same V2 computations as sensitivity to the suite of naturalistic statistics we use here.

Finally, although the results we present here provide evidence that sensitivity to the statistics of natural textures arises in V2 and that what sensitivity does exist within V1 results from feedback, we did not find major differences between laminae in V2. We have previously proposed a mechanism for how V2 neurons encode the statistical dependencies of naturalistic textures through computations analogous to those done by V1 complex cells (Freeman et al., 2013). Linear combinations of differently tuned V1 receptive fields are only weakly sensitive to higher-order correlations, but the addition of a rectifying and pooling stage that combines the output of multiple, different linear combinations can account for many features of V2 responses to naturalistic textures (Freeman et al., 2013: Ziemba 2016). Such an account is attractive since it casts the emergence of a novel form of selectivity in terms of known canonical computations. Given this framework, one might expect “V2 complex cells” sensitive to naturalistic image structure to predominate outside of the input layers of V2, analogous to the preponderance of traditional complex cells outside of the input layers of V1 (Hubel and Wiesel, 1968). We find only limited evidence for such a pattern, with the strongest V2 naturalness sensitivity found in the supragranular layers. This pattern is more prominent in single unit recordings in response to informative multipoint correlations (Yu et al., 2015). However, further work is needed to uncover the mechanisms by which naturalistic structure is extracted by V2 circuits.

## Acknowledgements

This work was supported by US National Institutes of Health grants EY04440, EY022428, and a National Science Foundation Graduate Research fellowship awarded to C.M.Z. The authors declare no competing financial interests.

